# The FlgN chaperone activates the Na^+^-driven engine of the flagellar protein export apparatus

**DOI:** 10.1101/2020.07.14.203299

**Authors:** Tohru Minamino, Miki Kinoshita, Yusuke V. Morimoto, Keiichi Namba

**Affiliations:** Graduate School of Frontier Biosciences, Osaka University, 1-3 Yamadaoka, Suita, Osaka 565-0871, Japan; Department of Physics and Information Technology, Faculty of Computer Science and Systems Engineering, Kyushu Institute of Technology, 680-4 Kawazu, Iizuka, Fukuoka 820-8502, Japan; RIKEN Spring-8 Center and Center for Biosystems Dynamics Research, 1-3 Yamadoaka, Suita, Osaka 565-0871, Japan; JEOL YOKOGUSHI Research Alliance Laboratories, Osaka University, 1-3 Yamadoaka, Suita, Osaka 565-0871, Japan

**Author notes:** Address correspondence to T. Minamino,.

## Abstract

The bacterial flagellar protein export machinery promotes H^+^-coupled translocation of flagellar proteins to the cell exterior. When the cytoplasmic ATPase complex does not function, the transmembrane export gate complex opens its Na^+^ channel and continues protein transport. However, it remains unknown how. Here we report that the FlgN chaperone acts as a switch to activate a backup export mechanism for the ATPase complex by activating the Na^+^-driven engine. Impaired interaction of FlhA with the FliJ subunit of the ATPase complex increased Na^+^-dependence of flagellar protein export. Deletion of FlgN inhibited protein export in the absence of the ATPase complex but not in its presence. Gain-of-function mutations in FlhA restored not only the FlgN defect but also the FliJ defect. We propose that the interaction of FlgN with FlhA opens the Na^+^ channel in the export engine, thereby maintaining the protein export activity in the absence of the active ATPase complex.

---

The bacterial flagellum is a macromolecular protein complex responsible for rapid and efficient movement of bacterial cells towards more suitable environments. The flagellum is composed of the basal body acting as a rotary motor, the hook as a universal joint and the filament as a helical propeller^1,2^. To construct the flagellum on the cell surface, a specialized protein export machinery is located at the flagellar base and transports flagellar structural subunits from the cytoplasm to the distal end of the growing flagellar structure. The flagellar export machinery is composed of a transmembrane export gate complex powered by proton motive force (PMF) across the cytoplasmic membrane and a cytoplasmic ATPase ring complex^3,4^. This export machinery is structurally and functionally similar to virulence-related type III secretion systems of pathogenic bacteria involved in direct injection of virulence effector proteins into eukaryotic host cells^5^.

The export gate complex is located inside the basal body MS ring and acts as a proton/protein antiporter to drive H^+^-coupled protein translocation across the cytoplasmic membrane^3,4^. FliP, FliQ and FliR form a polypeptide channel complex for the translocation of export substrates across the cytoplasmic membrane^6,7^. FlhB associates with the FliP/FliQ/FliR complex and is postulated to coordinate gate opening of the polypeptide channel^8^. FlhA associates not only with the FliP/FliQ/FliR complex but also with the MS ring^9^. Because FlhA promotes the transit of H^+^ and Na^+^ across the cytoplasmic membrane, it seems to act as an export engine of the export gate complex^10,11^. The C-terminal cytoplasmic domains of FlhA (FlhA_C_) and FlhB (FlhB_C_) project into the central cavity of the basal body C ring and form a docking platform for the cytoplasmic ATPase complex (FliH, FliI, FliJ), flagellar chaperones (FlgN, FliS, FliT) and export substrates^12,13^. This docking platform coordinates the order of flagellar protein export with assembly in a highly organized and well-controlled manner^14^.

FliH, FliI and FliJ form the cytoplasmic ATPase ring complex at the flagellar base^15^. This ATPase ring complex is not essential for flagellar protein export in *Salmonella enterica* serovar Typhimurium (hereafter referred to *Salmonella*)^16,17^ but ensures robust and efficient energy coupling of flagellar protein export^18,19^. The FliI ATPase hydrolyses ATP and activates the export gate complex through an interaction between FlhA_C_ and FliJ located at the center of the FliI hexamer ring, allowing the transmembrane export gate complex to become an active protein transporter to couple the H^+^ influx through the FlhA ion channel with the translocation of export substrates through the FliP/FliQ/FliR polypeptide channel^20–22^. Interestingly, the export gate complex also uses sodium motive force (SMF) across the cytoplasmic membrane to drive Na^+^-coupled protein export when the cytoplasmic ATPase ring complex does not function properly^10^. Because the FlhA ion channel conducts both H^+^ and Na^+^, there is a plausible hypothesis that the ATPase complex may switch the ion channel properties of FlhA from a dual ion channel mode to a highly efficient H^+^ channel through an interaction between FlhA_C_ and FliJ^10^. However, it remains unknown how it does.

To clarify this question, we analyzed the export properties of the *Salmonella* MM104-3 strain [*fliJ(*Δ*13*–*24) fliH(*Δ*96*–*97)*], in which the extragenic *fliH(*Δ*96*–*97)* suppressor mutation partially rescues the interaction of FliJ(Δ13–24) with FlhA_C_, thereby restoring flagellar formation in the presence of the *fliJ(*Δ*13-24)* mutation^20^. We show that the MM104-3 cells use SMF to efficiently produce flagella and that an interaction of FlhA_C_ with FlgN is essential for this Na^+^-coupled protein export. We will discuss a possible backup mechanism of the flagellar export engine.

## Results

### Effect of SMF on flagellar protein export by MM104-3 cells

Deletion of residues 13–24 of FliJ reduces the binding affinities of FliJ for FliI and FlhA_C_^20^. Deletion of residues 96–97 of FliH partially improves the interaction between FliJ(Δ13–24) and FlhA_C_, thereby restoring motility of the *fliJ(*Δ*13*–*24)* mutant to a considerable degree^20^. To clarify whether the impaired FliJ-FlhA_C_ interaction induces opening of the Na^+^ channel of FlhA, we analyzed the effect of SMF on flagellar formation by the wild-type and MM104-3 [*fliJ(*Δ*13*–*24) fliH(*Δ*96*–*97)*] cells. We used a condition of an external pH of 7.5 to diminish the chemical potential gradient of H^+^ (Ref. 23). Motility of the MM104-3 strain was better in the presence of 100 mM NaCl than in its absence (Fig. 1a). Consistently, the levels of FlgD and FliC secreted by the MM104-3 cells were much higher in the presence of 100 mM NaCl than in its absence (Fig. 1b), indicating that Na^+^ facilitates protein export by the MM104-3 cells. In contrast, both motility of and flagellar protein export by wild-type cells did not show a Na^+^ dependence (Fig. 1), in agreement with a previous report^10^.

**Fig. 1.**
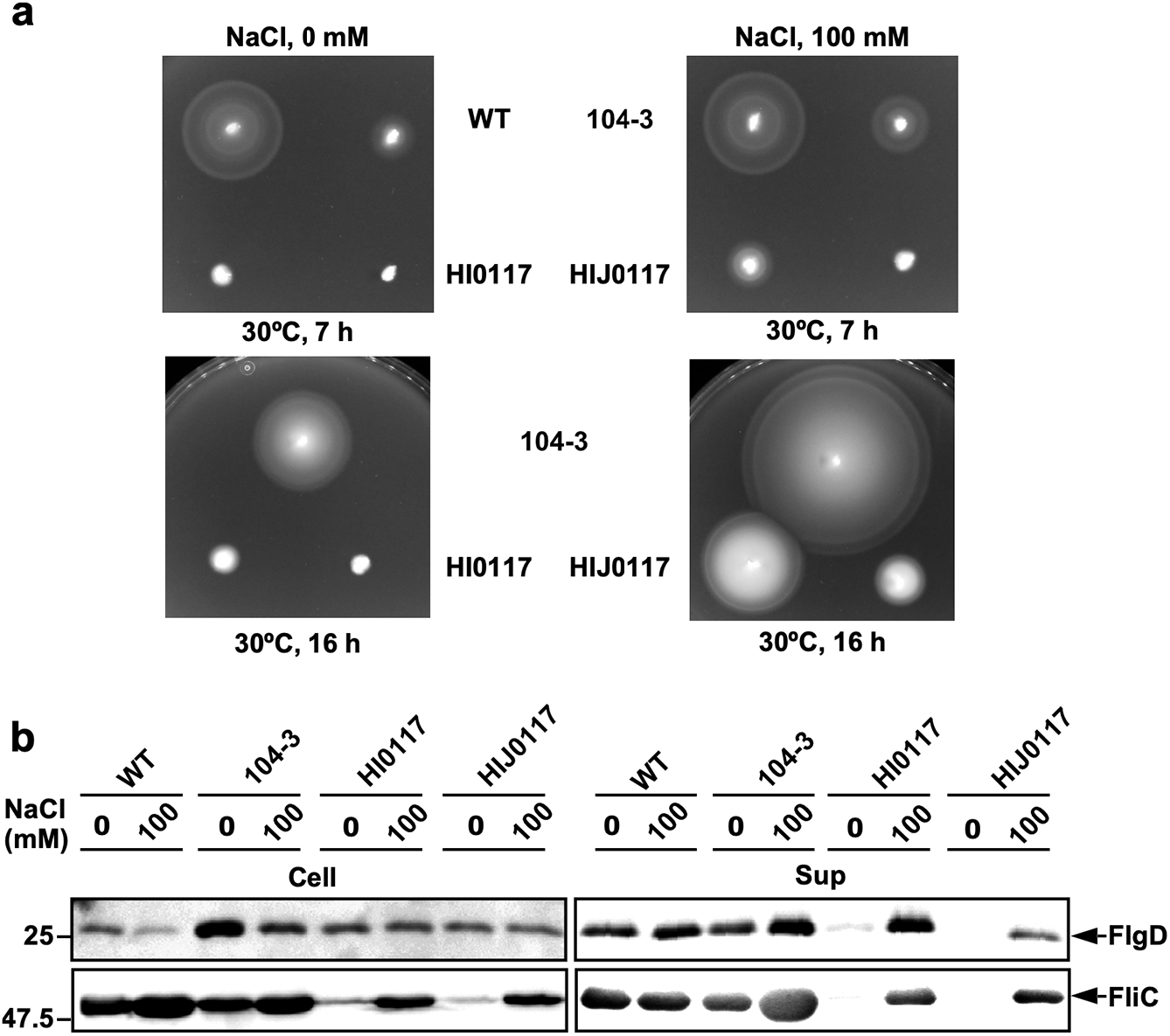
Effect of Na^+^ ions on flagellar protein export by MM104-3 cells. **(a)** Motility of SJW1103 (wild-type, indicated as WT), MM104-3 [*fliJ(*Δ*13-24) fliH(*Δ*96-97)*, indicated as 104-3], MMHI0117 [Δ*fliH-fliI flhB(P28T)*, indicated as HI0117] and MMHIJ0117 [Δ*fliH-fliI-fliJ flhB(P28T)*, indicated as HIJ0117] in 0.35% soft agar plates in the absence (left panels) and presence (right panels) of 100 mM NaCl. **(b)** Effect of Na^+^ on flagellar protein export at external pH 7.5. Immunoblotting, using polyclonal anti-FlgD or anti-FliC antibody, of whole cell proteins (Cell) and culture supernatant fractions (Sup) prepared from the above strains grown exponentially at 30°C in T-broth (pH 7.5) with or without 100 mM NaCl.

To quantitatively analyze the efficiency of flagellar assembly, we labelled the filaments with a fluorescent dye (Fig. 2a) and measured the number and length of the filaments (Supplementary Table 1). Wild-type cells produced the filaments with an average number of 2.7 ± 1.1 per cell (mean ± SD, n = 152) in the absence of NaCl and 3.3 ± 1.5 per cell (n = 153) in the presence of 100 mM NaCl (Fig. 2b). An average filament length was 11.3 ± 2.1 μm (n = 50) in the absence of NaCl and 12.9 ± 2.5 μm (n = 50) in the presence of 100 mM NaCl (Fig. 2c). Because the transcription levels of flagellar genes were not increased by adding 100 mM NaCl^10^, we suggest that the wild-type protein export apparatus also utilizes Na^+^ as a coupling ion to some extent. In the absence of NaCl, 38.5% of the MM104-3 cells had no filaments while the remaining population produced the filaments with an average number of 1.3 ± 0.5 per cell (n = 118) (Fig. 2b). The average filament length was 7.0 ± 2.9 μm (n = 50), which was about 1.6-fold shorter than the wild-type length in the absence of NaCl (Fig. 2c), indicating that the filament growth rate of the MM104-3 strain is slower than that of the wild type. In contrast, 87.3% of the MM104-3 cells produced the filaments in the presence of 100 mM NaCl with an average number of 2.0 ± 1.0 per cell (n = 145) (Fig. 2b). The average filament length was 10.7 ± 3.1 μm (n = 50), which was about 1.5-fold longer than that without NaCl (Fig. 2c). These results suggest that the transmembrane export gate complex of the MM104-3 strain actively uses SMF in addition to PMF to transport flagellar structural proteins during flagellar assembly. Therefore, we propose that the impaired interaction between FliJ and FlhA_C_ turns on a backup engine by allowing the export gate complex to use Na^+^ to drive flagellar protein export.

**Fig. 2.**
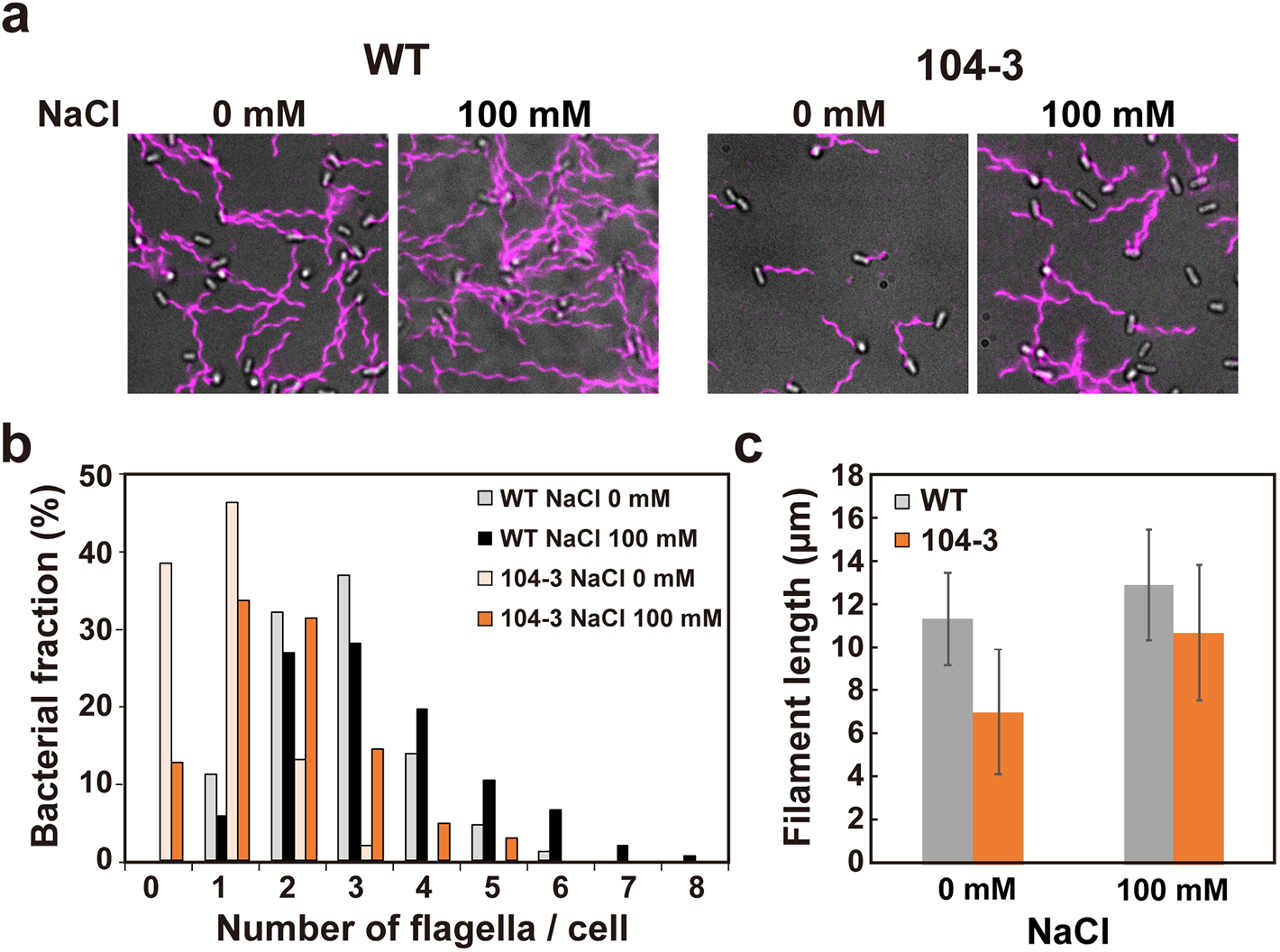
Measurements of the number and length of flagellar filaments produced by MM104-3 cells. **(a)** Fluorescent images of SJW1103 (WT) and MM104-3 [*fliJ(*Δ*13-24) fliH(*Δ*96-97)*, indicated as 104-3] grown in T-broth (pH 7.5) with or without 100 mM NaCl. Flagellar filaments were labelled with a fluorescent dye, Alexa Fluor 594. The fluorescence images of the filaments labelled with Alexa Fluor 594 (magenta) were merged with the bright field images of the cell bodies. **(b)** Distribution of the number of flagellar filaments in the SJW1103 and MM104-3 cells. **(c)** Measurements of the length of the flagellar filaments. Filament length is the average of 50 filaments, and vertical lines are standard deviations. (See Supplementary Table 1)

### Effect of FlgN deletion on flagellar protein export by MM104-3 cells

The *Salmonella* MMHI0117 strain [Δ*fliH-fliI flhB(P28T)*] is a pseudorevertant isolated from a mutant with deletion of FliH and FliI, which form the ATPase complex^16^. A non-functional FliJ variant, GST-FliJ, binds to FlhA and inhibits flagellar protein export by the MMHI0117 cell^24^, which produce a few flagella in the presence of 100 mM NaCl but not in its absence (Fig. 1)^10^. To identify which flagellar protein is responsible for the export engine to prefer SMF over PMF, GST-FliJ was over-expressed in the MMHI0117 cells as a bait, and then whole cell lysates were subjected to GST affinity chromatography. In addition to FlhA, FlgN co-purified with GST-FliJ but not with GST alone (Supplementary Fig. 1). FlgN is a flagellar export chaperone specific for two hook-filament junction proteins, FlgK and FlgL, and facilitates the docking of these two proteins to FlhA_C_ for rapid and efficient protein export^25–27^. FlgN binds to FlhA_C_ with nanomolar affinity even in the absence of FlgK and FlgL^28^, raising the possibility that an interaction between FlgN and FlhA_C_ could induce the opening of the Na^+^ channel of FlhA when FliJ does not function properly.

To clarify this possibility, we introduced a Δ*flgN*::*tetRA* allele into the MM104-3 strain by P22-mdeiated transduction and analyzed motility of the MM104-3 Δ*flgN*::*tetRA* cells in soft agar. Deletion of FlgN caused no motility of the MM104-3 cells (Fig. 3a). Consistently, no filaments were seen in the MM104-3 Δ*flgN*::*tetRA* mutant cells (Supplementary Fig. 2a). In contrast, about 35.2% of the Δ*flgN*::*tetRA* cells produced a single flagellar filament (Supplementary Fig. 2a), thereby generating a small motility ring on soft agar plates (Fig. 3a), in agreement with a previous report^29^.

**Fig. 3.**
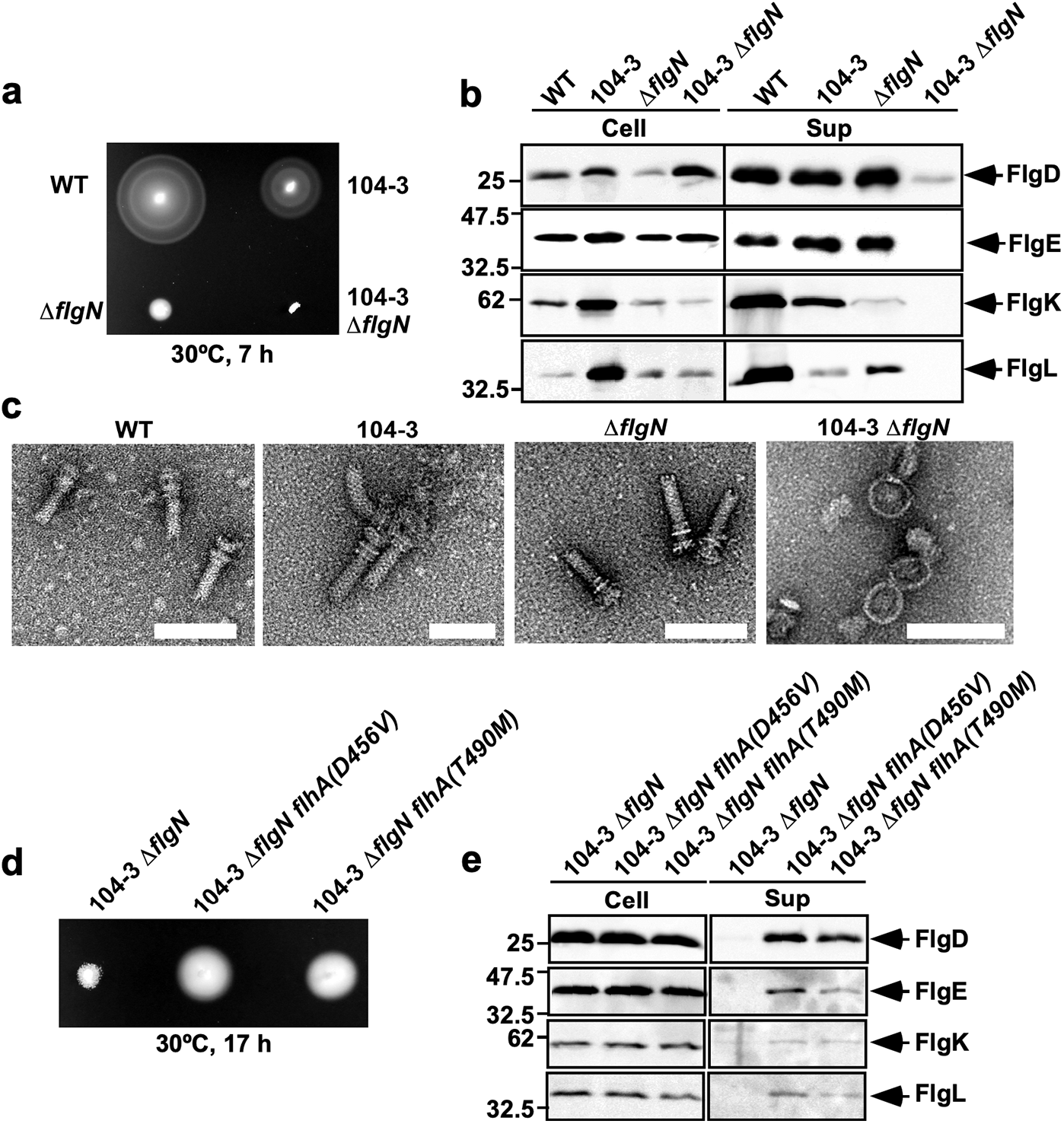
Effect of deletion of FlgN on flagellar protein export by MM104-3 cells. **(a)** Motility of SJW1103 (wild-type, indicated as WT), MM104-3 [*fliJ(*Δ*13-24) fliH(*Δ*96-97)*, indicated as 104-3], MM9001 (Δ*flgN*::*tetRA*, indicated as Δ*flgN*) and MM9003 [*fliJ(*Δ*13-24) fliH(*Δ*96-97)* Δ*flgN*::*tetRA*, indicated as 104-3 Δ*flgN*] in 0.35% soft agar plates containing 100 mM NaCl. **(b)** Immunoblotting, using polyclonal anti-FlgD (1st row), anti-FlgE (2nd row), anti-FlgK (3rd row) or anti-FlgL (4th row) antibody, of whole cell proteins (Cell) and culture supernatant fractions (Sup) prepared from the above strains. **(c)** Electron micrograms of hook-basal bodies isolated from the above stains. Scale bar shows 100 nm. **(d)** Motility of MM9003, MM9003-2 [*fliJ(*Δ*13-24) fliH(*Δ*96-97)* Δ*flgN*::*tetRA flhA(D456V)*, indicated as 104-3 Δ*flgN flhA(D456V)*] and MM9003-3 [*fliJ(*Δ*13-24) fliH(*Δ*96-97)* Δ*flgN*::*tetRA flhA(T490M)*, indicated as 104-3 Δ*flgN flhA(T490M)*] in 0.35% soft agar plates containing 100 mM NaCl. **(e)** Immunoblotting, using polyclonal anti-FlgD (1st row), anti-FlgE (2nd row), anti-FlgK (3rd row) or anti-FlgL (4th row) antibody, of whole cell proteins (Cell) and culture supernatant fractions (Sup) prepared from the above strains.

We next analyzed the impact of depletion of FlgN on flagellar protein export by the MM104-3 strain. A loss of FlgN considerably reduced the levels of FlgK and FlgL secreted by wild-type cells but not those of the hook-capping protein, FlgD and the hook protein, FlgE (Fig. 3b). Consistently, the Δ*flgN*::*tetRA* cells produced hook-basal bodies (HBBs) without filament attached (Fig. 3c), in agreement with a previous report^29^. In contrast, FlgN deletion inhibited also the secretion of FlgD and FlgE by the MM104-3 cells (Fig. 3b). The MM104-3 cells produced HBBs whereas the MM104-3 Δ*flgN*::*tetRA* cells produced only the MS-C ring structures (Fig. 3c). When FlgN was expressed from a pTrc99A based plasmid in the MM104-3 Δ*flgN*::*tetRA* cells, the motility was restored to a level comparable to that of the MM104-3 strain (Supplementary Fig. 2b). These results indicate that FlgN is essential for the export of flagellar structural proteins by the MM104-3 cells.

### Isolation of pseudorevertants from the MM104-3 Δ*flgN*::*tetRA* strain

It has been shown that the *flhA(D456V)* and *flhA(T490M)* mutations partially restore the FlgN defect^26^. Therefore, we introduced the *flhA(D456V)* and *flhA(T490M)* alleles into the MM104-3 Δ*flgN*::*tetRA* cells by P22-mdeiated transduction to see whether these *flhA* mutations rescue the protein export activity of the MM104-3 Δ*flgN*::*tetRA* cells. The *flhA(D456V)* and *flhA(T490M)* mutations improved motility of and flagellar protein export by the MM104-3 Δ*flgN*::*tetRA* cells (Fig. 3d and 3e), suggesting that the *flhA(D456V)* and *flhA(T490M)* mutations allow FlhA_C_ to adopt a certain conformation mimicking a FlgN-bound state of FlhA_C_ even in the absence of FlgN. Because FlgN alone binds to FlhA_C_ with nanomolar affinity^28^, we suggest that the interaction between FlgN and FlhA_C_ is critical for the export of FlgD and FlgE by the MM104-3 strain. Furthermore, to understand why the MM104-3 strain requires FlgN for the export of FlgD and FlgE, we isolated five pseudorevertants from the MM104-3 Δ*flgN*::*tetRA* strain. Motility of these pseudorevertants was better than that of the parent strain but much poorer than that of the MM104-3 strain (Supplementary Fig. 3a). The secretion levels of FlgD, FlgE and FliC by these pseudorevertants were almost the same as those by the MM104-3 cells (Supplementary Fig. 3b). However, the secretion level of FlgK by these pseudorevertants was not restored (Supplementary Fig. 3b, 3rd row) because they lacked FlgN. Therefore, we propose that FlgN acts not only as an export chaperone specific for FlgK and FlgL but also as a switch to activate a backup export mechanism for the cytoplasmic ATPase complex in the MM104-3 cells by activating the Na^+^-coupled protein export engine.

DNA sequence analysis established that one of the suppressor mutations was the L244R missense mutation in the ATPase domain of FliI (seen twice) (Supplementary Fig. 3c). Leu-244 makes hydrophobic contacts with Val-70, Glu-283 and Ile-284, and so the *fliI(L244R)* mutation is likely to affect the FliJ-FliI interface. The secretion of FlgD by the MM104-3 Δ*flgN*::*tetRA flhA(D456V)* and MM104-3 Δ*flgN*::*tetRA fliI(L244R)* cells still showed a Na^+^-dependence in a way similar to the MM104-3 strain (Supplementary Fig. 3d). Interestingly, the *flhA(D456V)* and *fliI(L244R)* mutations increased the secretion level of FlgD even in the absence of Na^+^ (Supplementary Fig. 3d). Because it has been shown that the *fliI(L244R)* mutation partially restores the interaction of FliJ(Δ13–24) with FlhA_C_^20^, we suggest that ATP hydrolysis by FliI with the L244R mutation may allow FliJ(Δ13–24) to induce conformational rearrangements of the export gate complex through the FliJ-FlhA_C_ interaction even in the absence of FlgN, thereby allowing the gate complex to become an active export engine to utilize both H^+^ and Na^+^ to drive flagellar protein export.

The other suppressor mutations were located within the *flgM* gene, which encodes a transcription regulator for the class 3 flagellar proteins, such as FliC. One caused a stop codon (TAA) at a position of Ser-7 of FlgM, resulting in truncation of the C-terminal region of FlgM, and the others were large deletions. A loss-of-function of FlgM results in a considerable increment in the transcription levels of flagellar genes^30^. Consistently, the cellular levels of FlgK and FliC seen in the MM104-3 Δ*flgN*::*tetRA* Δ*flgM* cells were larger than those in its parental strain (Supplementary Fig. 3b, 3rd and 4th rows). Therefore, we suggest that these *flgM* mutations considerably increase the cytoplasmic levels of both export substrates and cytoplasmic export apparatus components, allowing the MM104-3 Δ*flgN*::*tetRA* strain to export flagellar proteins to a considerable degree.

### Effect of deletion of FliJ residues 13–24 on the interactions of FlgN and FlhA_C_

FliJ binds to FlgN with a K_D_ of 22 μM^31^, raising the possibility that FlgN acts as a backup for FliH and FliI to support efficient docking of FliJ to FlhA_C_. To clarify this possibility, we first investigated whether deletion of residue 13–24 of FliJ affects the interaction of FliJ with FlgN by GST affinity chromatography. FlgN co-purified with GST-FliJ(Δ13–24) (Supplementary Fig. 4a), indicating that this deletion does not abolish the FliJ-FlgN interaction. To assess the strength of the FliJ-FlgN interaction, we determined the kinetic parameters for the binding of FlgN to immobilized GST-FliJ or GST-FliJ(Δ13–24) by surface plasmon resonance (SPR). Steady state analysis of SPR data with a 1 to 1 binding model indicated that the K_D_ values for the FliJ-FlgN and FliJ(Δ13–24)-FlgN interactions were 27.4 ± 13.0 μM and 7.74 ± 0.12 μM, respectively (Supplementary Fig. 4b), indicating that the deletion increases the binding affinity of FliJ for FlgN by about 3.5-fold. We next investigated whether FlgN supports FliJ(Δ13–24) to interact with FlhA_C_. The amounts of FlhA_C_ co-purified with GST-FliJ(Δ13–24) were much lower than those with GST-FliJ (Supplementary Fig. 4c and 4d, upper panels), in agreement with a previous report^20^. FlgN did not improve the interaction of FliJ(Δ13–24) with FlhA_C_ (lower panels). These results suggest that a direct interaction between FlgN and FlhA_C_ turns on Na^+^-coupled protein export by the MM104-3 strain when the FliJ-FlhA_C_ interaction is compromised.

### Effect of the *flhA(D456V)* and *flhA(T490M)* mutations on Na^+^-coupled protein export by cells lacking FliH, FliI and FliJ

It has been shown that FliJ is more important for Na^+^-coupled flagellar protein export than FliH and FliI^20^ because the protein export activity of the Δ*fliH-fliI-fliJ flhB(P28T)* strain (MMHIJ0117) is much lower than that of the MMHI0117 strain (Fig. 1). To confirm this, we first analyzed the number and length of flagellar filaments produced by the MMHIJ0117 cells (Fig. 4a and Supplementary Table 2). About 78.2% of the MMHI0117 cells produced the filaments with an average number of 1.6 ± 0.7 per cell (n = 266) and an average length of 7.8 ± 2.5 μm (n = 50) (Fig. 4b and c), in agreement with a previous report^19^. In contrast, only 13.5% of the MMHIJ0117 cells produced the filaments with an average number of 1.1 ± 0.2 per cell (n = 57) and an average length of 5.1 ± 2.2 μm (n = 50) (Fig. 4b and c). Therefore, we conclude that FliJ acts as a more important activator of the transmembrane export gate complex than FliH and FliI.

**Fig. 4.**
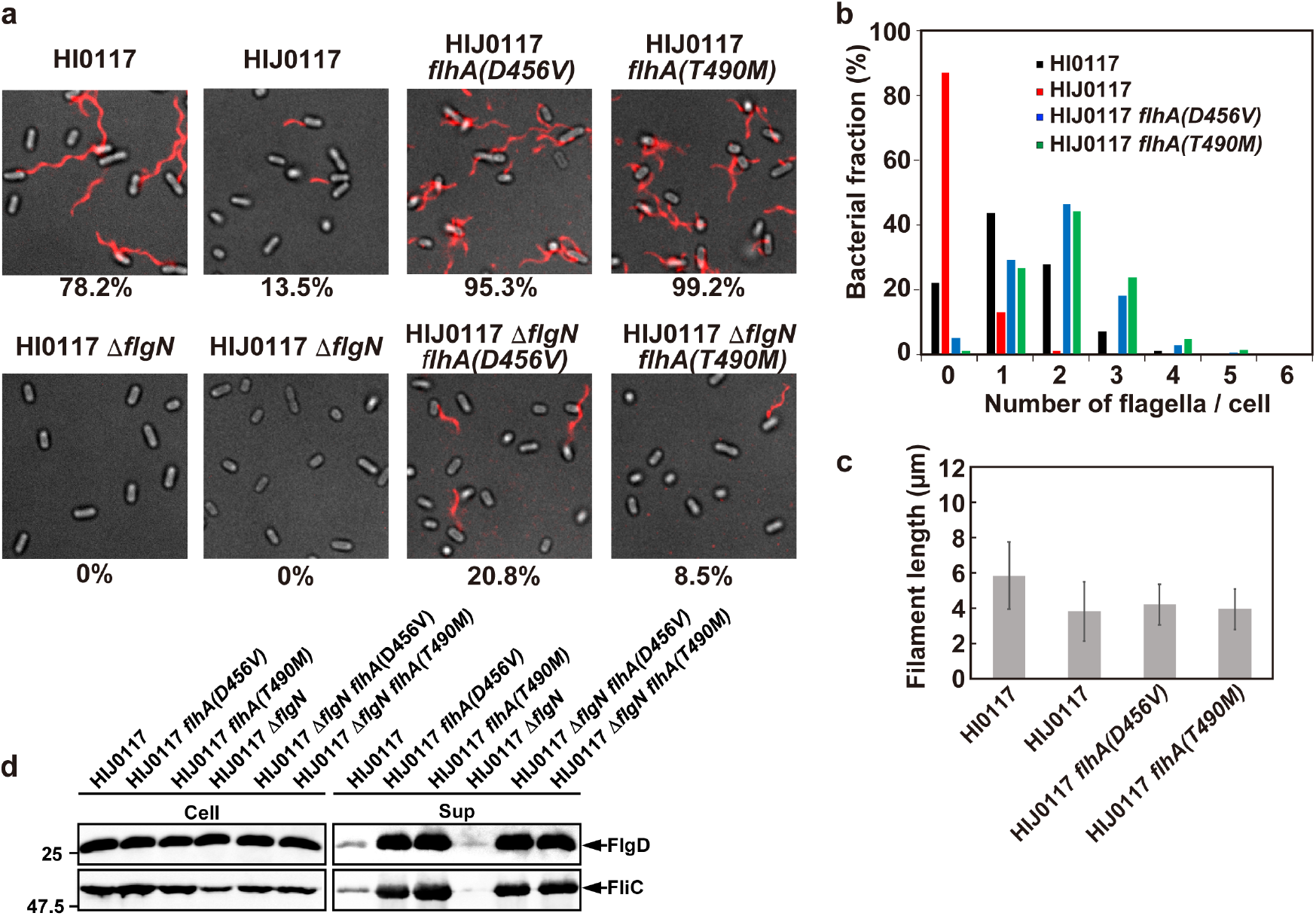
Effects of gain-of-function mutations in FlhA and deletion of FlgN on flagellar protein export and assembly by MMHIJ0117 cells. **(a)** Fluorescent images of MMHI0117 [Δ*fliH-fliI flhB(P28T)*, indicated as HI0117], MMHIJ0117 [Δ*fliH-fliI-fliJ flhB(P28T)*, indicated as HIJ0117], MMHIJ0117-2 [Δ*fliH-fliI-fliJ flhB(P28T) flhA(D456V)*, indicated as HIJ0117 *flhA(D456V)*], MMHIJ0117-3 [Δ*fliH-fliI-fliJ flhB(P28T) flhA(T490M)*, indicated as HIJ0117 *flhA(T490M)*], MM9002 [Δ*fliH-fliI flhB(P28T)* Δ*flgN*::*tetRA*, indicated as HI0117 Δ*flgN*], MM9004 [Δ*fliH-fliI-fliJ flhB(P28T)* Δ*flgN*::*tetRA*, indicated as HIJ0117 Δ*flgN*], MM9004-2 [Δ*fliH-fliI-fliJ flhB(P28T)* Δ*flgN*::*tetRA flhA(D456V)*, indicated as HIJ0117 Δ*flgN flhA(D456V)*] and MM9004-3 [Δ*fliH-fliI-fliJ flhB(P28T)* Δ*flgN*::*tetRA flhA(T490M)*, indicated as HIJ0117 Δ*flgN flhA(T490M)*] grown in T-broth (pH 7.5) containing 100 mM NaCl. Flagellar filaments were labelled with Alexa Fluor 594. The fluorescence images of the filaments labelled with Alexa Fluor 594 (red) were merged with the bright field images of the cell bodies. **(b)** Distribution of the number of flagellar filaments in the MMHI0117, MMHIJ0117, MMHIJ0117-2 and MMHIJ0117-3 cells. **(c)** Measurements of the length of the flagellar filaments. Filament length is the average of 50 filaments, and vertical lines are standard deviations. (See Supplementary Table 2) **(d)** Immunoblotting, using polyclonal anti-FlgD (1st row) or anti-FliC (2nd row) antibody, of whole cell proteins (Cell) and culture supernatant fractions (Sup) prepared from the above strains.

To investigate whether FlgN becomes essential in the MMHIJ0117 strain in a way similar to the MM104-3 strain, we introduced a Δ*flgN*::*tetRA* allele into the MMHIJ0117 strain and found that FlgN deletion inhibited the secretion of FlgD and FliC by the MMHIJ0117 strain (Fig. 4d), thereby conferring a Fla^−^ phenotype (Fig. 4a). When FlgN was expressed from a plasmid, the motility was recovered to a level comparable to that of the MMHIJ0117 strain (Supplementary Fig. 2b). Because the secretion of FlgD and FliC by the MMHIJ0117 cells showed a clear Na^+^-dependence (Fig. 1b), these results suggest that FlgN is more important for Na^+^-coupled flagellar protein export than FliJ when FliH and FliI are missing.

The *flhA(D456V)* and *flhA(T490M)* mutations have been isolated as gain-of-function mutations to restore motility of the MMHI0117 Δ*flgN*::tetRA strain. Therefore, we re-examined whether the loss of FlgN inhibits flagellar protein export by the MMHI0117 strain. No FlgD and FliC were detected in the culture supernatant of the MMHI0117 Δ*flgN*::tetRA strain but the *flhA(D456V)* and *flhA(T490M)* mutations increased the secretion levels of FlgD and FliC by this mutant cells (Supplementary Fig. 5). Therefore, we conclude that the interaction of FlgN with FlhA_C_ becomes essential for Na^+^-coupled flagellar protein export by the export gate complex when the cytoplasmic ATPase complex is absent.

It has been shown that the *flhA(D456V)* and *flhA(T490M)* mutations enhance flagellar protein export by the MMHI0117 cells^19^, raising the possibility that these two *flhA* mutations can bypass the FliJ defect as well as the FlgN defect. To clarify this, we introduced these *flhA* alleles into the MMHIJ0117 strain and found that the *flhA(D456V)* and *flhA(T490M)* mutations increased the secretion levels of FlgD and FliC by more than 10-fold (Fig. 4d). Consistently, more than 95% of the MMHIJ0117 *flhA(D456V)* and MMHIJ0117 *flhA(T490M)* cells had a couple of flagellar filaments (Fig. 4a and b and Supplementary Table 2) although their average filament length was almost the same as that of the MMHIJ0117 strain (Fig. 4c). The loss of FlgN did not significantly reduce the secretion levels of FlgD and FliC by the MMHIJ0117 *flhA(D456V)* and MMHIJ0117 *flhA(T490M)* cells (Fig. 4d). Furthermore, 20.8% of the MMHIJ0117 Δ*flgN*::tetRA *flhA(D456V)* cells and 8.5% of the MMHIJ0117 Δ*flgN*::tetRA *flhA(T490M)* cells produced a single flagellar filament (Fig. 4a). Therefore, we conclude that the *flhA(D456V)* and *flhA(T490M)* mutations can activate the export gate complex to drive Na^+^-coupled flagellar protein export in the absence of FliH, FliI, FliJ and FlgN.

## Discussion

The flagellar protein export machinery maintains its protein export activity against various internal and external perturbations, and to do so, this export machinery has evolved to become a dual fuel export engine to exploit both H^+^ and Na^+^ as the coupling ion rather than becoming an exclusive PMF-or SMF-dependent export engine^10,18,20^. The wild-type export engine predominantly uses H^+^ as a coupling ion. However, when the ATPase complex does not work properly, the export engine opens its Na^+^ channel to continue the assembly of flagella on the cell surface^10,20,22^, but it remained unknown how. Here we showed that an impaired interaction between FliJ and FlhA_C_ by the lack of ATPase activity activated the Na^+^-driven export engine to promote Na^+^-coupled protein export (Figs 1 and 2). We also found that an interaction between FlgN and FlhA_C_ became essential for Na^+^-coupled protein export (Figs 3 and 4). FlgN is an export chaperone specific for FlgK and FlgL^25^. FlgN facilitates the docking of FlgK and FlgL to the FlhA_C_ platform of the export gate complex for rapid and efficient protein export^26–28,31,32^. So, the loss of FlgN reduces the secretion levels of FlgK and FlgL, resulting in a considerable reduction in the probability of filament formation at the tip of the HBB^29^. The deletion of FlgN inhibited not only the export of FlgK and FlgL but also that of FlgD and FlgE by the MM104-3 cells (Fig. 3b). A gain-of-function mutation in FliI restored the export of FlgD and FlgE but not that of FlgK and FlgL (Supplementary Fig. 3). Because the wild-type export engine also used SMF to form flagellar filaments to some degree (Fig. 2b,c), we propose that FlgN acts not only as a substrate-specific export chaperone but also as a switch to activate a backup mechanism for the cytoplasmic ATPase complex by turning on the Na^+^-driven export engine through an interaction between FlgN and FlhA_C_ to compensate the occasional loss or inactivation of the ATPase complex during flagellar assembly (Fig 5).

**Fig. 5.**
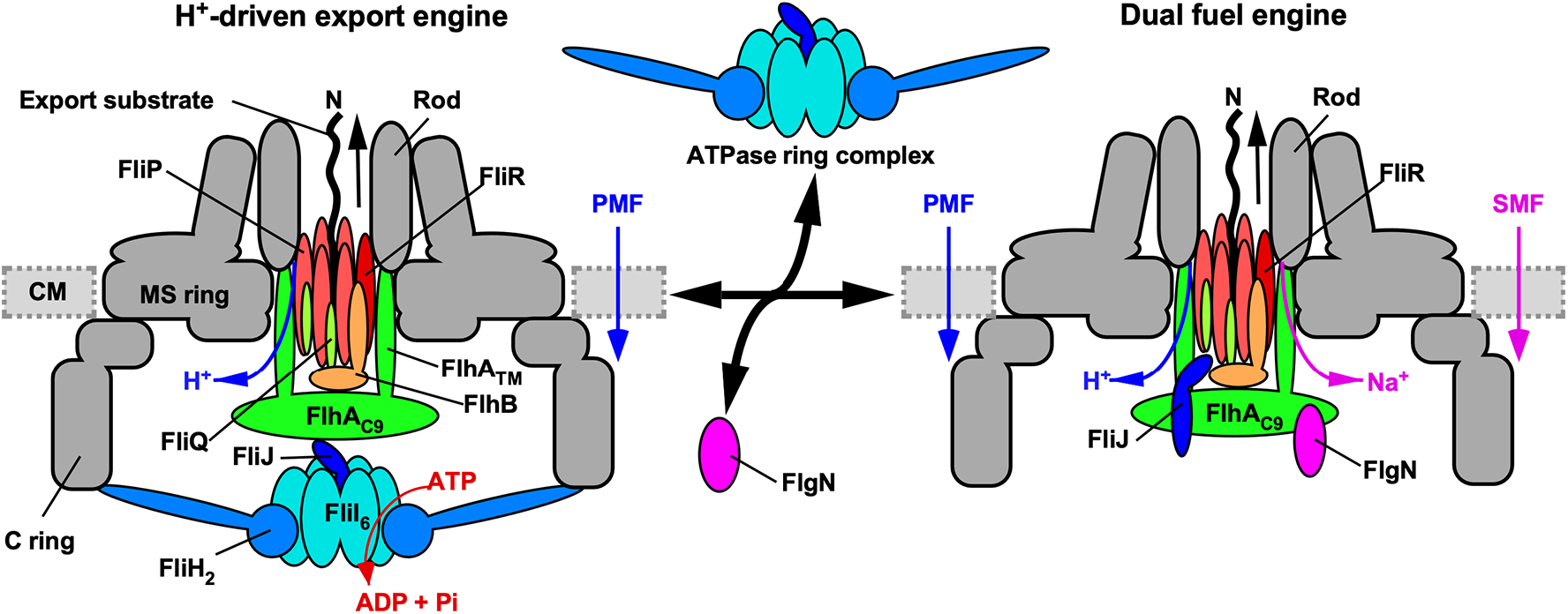
Schematic diagram of the flagellar protein export machinery. The flagellar protein export machinery is composed of a transmembrane export gate complex made of FlhA, FlhB, FliP, FliQ and FliR and a cytoplasmic ATPase complex consisting of FliH, FliI and FliJ. The export gate complex is located inside the MS ring and utilizes proton motive force (PMF) across the cytoplasmic membrane (CM) to drive proton (H^+^)-coupled flagellar protein export. FliP, FliQ and FliR form a polypeptide channel. FlhB associates with the FliP/FliQ/FliR complex and controls gate opening of the polypeptide channel. The C-terminal cytoplasmic domain of FlhA (FlhA_C_) projects into the central cavity of the C ring. The N-terminal transmembrane domain (FlhA_TM_) forms an ion channel for the translocation of H^+^ and sodium ion (Na^+^) from the periplasm to the cytoplasm. The cytoplasmic ATPase ring complex associates with the C ring through an interaction between FliH and a C ring protein, FliN. ATP hydrolysis by the FliI ATPase activates the export gate complex through an interaction between FliJ and FlhA_L_ connecting FlhA_C_ to FlhATM, becoming an active protein transporter to couple the H^+^ flow through the FlhA channel to the translocation of export substrates into the polypeptide channel. When the cytoplasmic ATPase complex does not function properly, FlgN binds to FlhA_C_ to open the Na^+^ channel of FlhA_TM_, allowing the export gate complex to utilize sodium motive force (SMF) across the cytoplasmic membrane to drive Na^+^-coupled protein export.

FlgN binds to a well-conserved hydrophobic dimple of FlhA_C_ including Asp-456, Phe-459 and Thr-490^26,33^. FliJ binds to the flexible linker region of FlhA (FlhA_L_) connecting FlhA_C_ to the N-terminal transmembrane domain, which forms an ion channel, and this FliJ-FlhA_L_ interaction activates the export gate complex to become an active H^+^-driven export engine^20,34^ (Fig 5). Consistently, the deletion of residues 328–351 of FlhA_L_ significantly weakened the FliJ-FlhA_C_ interaction but not the FlgN-FlhA_C_ interaction (Supplementary Fig. 6a,b). FlgN bound to FliJ(Δ13–24) but did not restore the impaired interaction between FliJ(Δ13–24) and FlhA_C_ at all (Supplementary Fig. 4). It has been shown that FliJ binds not only to FlgN^31^ but also to the FlgN/FlgK complex^26^. It has been shown that the FlgN/FlgK/FliJ trimeric complex docks to the FlhA_C_ platform (Supplementary Fig. 6c)^26^. When GST-FlgN/FlgK/FliJ complex was mixed with FlhA_C38K_ lacking residues 328–351 of FlhA_L_, only a very small amount of FliJ co-purified with GST-FlgN, FlgK and FlhA_C38K_ (Supplementary Fig. 6c), indicating that FliJ dissociates from FlgN upon binding of the FlgN/FlgK/FliJ complex to FlhA_C38K_. Because the protein transport activity of the transmembrane export gate complex was much higher in the presence of FliJ than in its absence (Fig. 1), we propose that the cytoplasmic FlgN/FliJ complex docks to the FlhA_C_ platform through an interaction between FlgN and FlhA_C_, thereby inducing the dissociation of the FlgN/FliJ complex into FlgN and FliJ subunits to bind to the hydrophobic dimple of FlhA_C_ and FlhA_L_, respectively, to fully activate the Na^+^-driven engine of the export gate complex in the absence of an active ATPase complex (Fig. 5).

A subpopulation of planktonic cells can rapidly move in the biofilm structure by rotating flagella to keep cells in the biofilm alive and healthy^35^. The second messenger molecule 3’-5’ cyclic diguanylate monophosphate not only inhibits the transcription of flagellar genes but also binds to the FliI ATPase to suppress flagellar assembly of the cells in the biofilm^36^. Total PMF is quite low in the cells living in the biofilm structure because the membrane voltage is quite small^37^. Therefore, it is likely that the flagellar protein export apparatus maintains the Na^+^-driven backup engine throughout the evolutionary process to efficiently generate flagellated cells in the biofilm structure.

## Methods

### Bacterial strains and plasmids

*Salmonella* strains and plasmids used in this study are listed in Supplementary Table 3.

### Motility assays in soft agar

Fresh colonies were inoculated onto soft agar plates [1% (w/v) tryptone, 0.35% Bacto agar] with or without 100 mM NaCl and incubated at 30°C. At least seven independent measurements were performed.

### Secretion assay

*Salmonella* cells were grown overnight in T-broth [1%(w/v) Bacto tryptone, 10 mM potassium phosphate pH 7.5] without 100 mM NaCl. A 50 μl of the overnight culture was inoculated into a 5 ml of fresh T-broth with or without 100 mM NaCl and incubated at 30 °C with shaking until the cell density had reached an OD_600_ of ca. 1.4–1.6. Cultures were centrifuged to obtain cell pellets and culture supernatants. The cell pellets were resuspended in a SDS-loading buffer (62.5 mM Tris-HCl, pH 6.8, 2% SDS, 10% glycerol, 0.001% bromophenol blue) containing 1 μl of 2-mercaptoethanol, normalized to a cell density to give a constant number of cells. Proteins in the culture supernatants were precipitated by 10% trichloroacetic acid, suspended in the Tris/SDS loading buffer (one volume of 1 M Tris, nine volumes of 1 X SDS loading buffer) containing 1 μl of 2-mercaptoethanol and heated at 95°C for 3 min. After Sodium Dodecyl Sulfate–polyacrylamide gel electrophoresis (SDS–PAGE), immunoblotting with polyclonal anti-FlgD, anti-FlgE, anti-FlgK, anti-FlgL or anti-FliC antibody was carried out as described previously^38^. Detection was performed with an ECL plus immunoblotting detection kit (GE Healthcare). Chemiluminescence signals were captured by a Luminoimage analyzer LAS-3000 (GE Healthcare). All image data were processed with Photoshop software CS6 (Adobe). At least three independent experiments were performed.

### Observation of flagellar filaments with a fluorescent dye

The flagellar filaments produced by *Salmonella* cells were labelled using anti-FliC antibody and anti-rabbit IgG conjugated with Alexa Fluor 594 (Invitrogen) as described previously^21^. The cells were observed by fluorescence microscopy as described previously^39^. Fluorescence images were analyzed using ImageJ software version 1.52 (National Institutes of Health).

### Preparation of HBBs

HBBs and MS-C ring complexes were prepared as described before^19^. Samples were negatively stained with 2% uranyl acetate. Electron micrographs were recorded with a JEM-1011 transmission electron microscope (JEOL, Tokyo, Japan) operated at 100 kV and equipped with a F415 CCD camera (TVIPS, Gauting, Germany) at a magnification of x5,500, which corresponds to 2.75 nm per pixel.

### Pull-down assays by GST chromatography

His-FlhA_C_ and His-FlgN were purified by Ni affinity chromatography with a nickel-nitriloacetic acid (Ni-NTA) agarose column (QIAGEN) as described previously^12^. Fractions containing His-FlhA_C_ or His-FlgN were dialyzed overnight against PBS (8 g of NaCl, 0.2 g of KCl, 3.63 g of Na_2_HPO_4_•12H_2_O, 0.24 g of KH_2_PO_4_, pH 7.4 per liter) at 4°C. GST-FliJ, GST-FliJ(Δ13–24) and GST-FlgN were purified by GST affinity chromatography using a glutathione Sepharose 4B column (GE Healthcare) as described previously^28^. Fractions containing GST-tagged proteins were pooled and dialyzed overnight against PBS at 4°C with three changes of PBS. Detail protocols for pull-down assays by GST affinity chromatography have been described previously^26,28^. At least three independent experiments were carried out.

### Surface plasmon resonance (SPR)

SPR measurements were carried out at 25°C using a Biacore X instrument (GE Healthcare) as described previously^28^. The K_D_ value was calculated as described in the manufacturer’s instructions (GE Healthcare).

## Acknowledgements

We acknowledge Yumi Inoue and Yasuyo Abe for technical assistance. This work was supported in part by JSPS KAKENHI Grant Numbers JP26293097 and JP19H03182 (to T.M.), JP18K14638 and JP20K15749 (to M.K.), JP15H05593 and JP18K06159 (to Y.V.M) and JP25000013 (to K.N.) and MEXT KAKENHI Grant Numbers JP15H01640 (to T.M.) and JP26115720 and JP15H01335 (to Y.V.M). This work has also been partially supported by JEOL YOKOGUSHI Research Alliance Laboratories of Osaka University to K.N.

## Author Contributions

T.M. and K.N. conceived and designed research; T.M., M.K. and Y.V.M. performed experiments; T.M., M.K. and Y.V.M. analyzed the data, and T.M. and K.N. wrote the paper based on discussion with other authors.

## Materials & Correspondence

Correspondence and requests for materials should be addressed to T.M.

