## Supplementary information for "The FlgN chaperone activates the Na^+^-driven engine of the flagellar protein export apparatus"

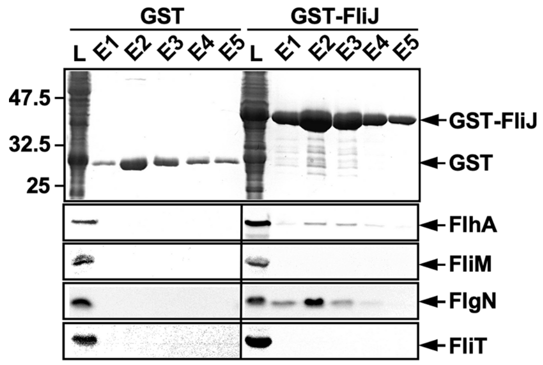

**Supplementary Fig. 1. Pull-down assays by GST affinity chromatography.** Cell lysates (indicated as L) prepared from *Salmonella* MMHI0117 [(*ΔfliH-fliI flhB(P28T)*] cells expressing GST or GST-FliJ were loaded onto a GST column. After washing with 15 ml of PBS, proteins were eluted with 5 ml of 50 mM Tris-HCl, pH 8.0, 10 mM reduced glutathione. Elution fractions were analysed by both CBB staining (1st row) and immunoblotting with anti-$FlhA_C$ (2nd row), anti-FliM (3rd row), anti-FlgN (4th row) or anti-FliT (5th row) because it has been reported that FliJ binds to $FlhA_C$, FliM, FlgN and FliT *in vitro*.

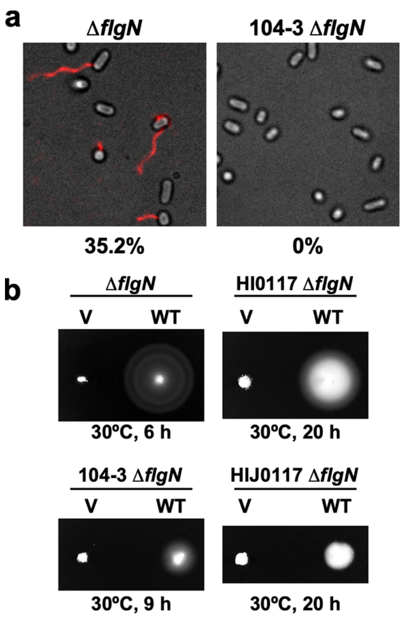

**Supplementary Fig. 2. Effect of FlgN deletion on flagellar filament formation. (a)** Fluorescent images of MM9001 (Δ*flgN*::*tetRA*, indicated as Δ*flgN*) and MM9003 (MM104-3 Δ*flgN*::*tetRA*, indicated as 104-3 Δ*flgN*). The flagellar filaments were labeled with Alexa Fluor 594. The fluorescence images of the filaments labeled with Alexa Fluor 594 (red) were merged with the bright field images of the cell bodies. About 35.2% of the MM9001 cells produced a single flagellar filament, but the MM9003 cells produced no filaments. **(b)** Motility of MM9001, MM9002 (MMHI0117 Δ*flgN*::*tetRA*, indicated as HI0117 Δ*flgN*), MM9003 or MM9004 (MMHIJ0117 Δ*flgN*::*tetRA*, indicated as HIJ0117 Δ*flgN*) harbouring pTrc99A (V) or pMMGN140 (FlgN) in soft agar.

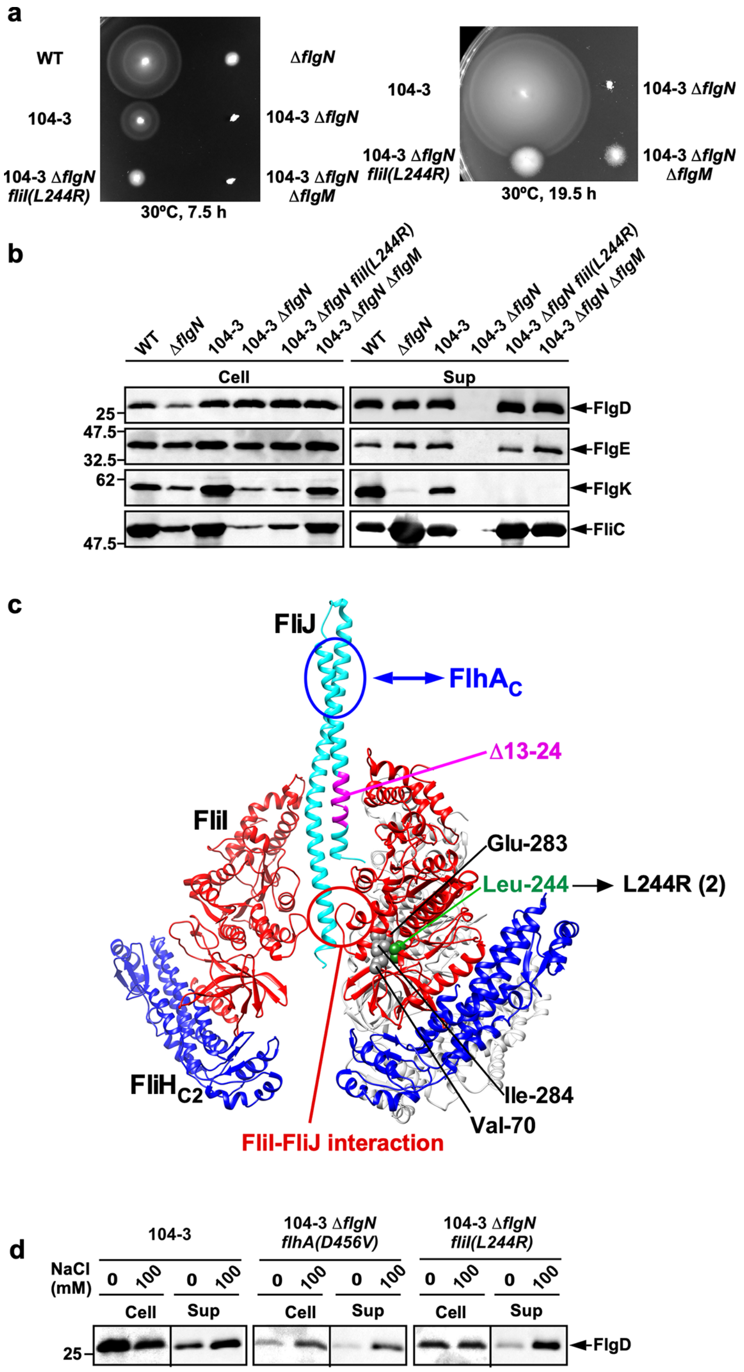

**Supplementary Fig. 3. Isolation of gain-of-function mutants from the MM104-3 Δ*flgN* strain.** **(a)** Motility of SJW1103, MM104-3, MM9001 (Δ*flgN*), MM9003 (MM104-3 Δ*flgN*), MM9003-4 [MM104-3 Δ*flgN fliI(L244R)*] and MM9003-5 [MM104-3 Δ*flgN flgM(S7ochre)*] in soft agar. Plates were incubated at 30ºC **(b)** Secretion of FlgD, FlgE, FlgK and FliC. Immunoblotting, using polyclonal anti-FlgD (1st row), anti-FlgE (2nd row), anti-FlgK (3rd row) or anti-FliC (4th row) antibody, of whole cell proteins (Cell) and culture supernatant fractions (Sup) prepared from the above strains. **(c)** Location of a gain-of-function mutation in FliI. Atomic model of the cytoplasmic ATPase ring complex consisting of FliH, FliI and FliJ. Cα backbone traces of the FliH$_{C2}$-FliI complex (PDB ID: 5B0O) and FliJ (PDB ID: 3AJW) are shown. The FliH$_{C2}$, FliI and FliJ subunits are colored blue, red, and cyan, respectively. Residues 13-24 of

FliJ are shown in magenta. Leu-244 makes hydrophobic contacts with Val-70, Glu-283 and Ile-284, and so the *fliI(L244R)* mutation, which restores motility of the MM104-3 Δ*flgN* cells to a significant degree, presumably affects an interface between FliI and FliJ. A well-conserved surface of FliJ is involved in the interaction with $FlhA_C$. **(d)** Effect of $Na^+$ on FlgD secretion by gain-of-function mutants. Immunoblotting, using polyclonal anti-FlgD antibody, of whole cell proteins (Cells) and culture supernatant fractions (Sup) prepared from the MM104-3, MM9003-2 [MM104-3 Δ*flgN flhA(D456V)*] (middle panels) and MM9003-4 [MM104-3 Δ*flgN fliI(L244R)*] cells (right panels) grown exponentially at 30ºC in T-broth with or without 100 mM NaCl at an external pH of 7.5.

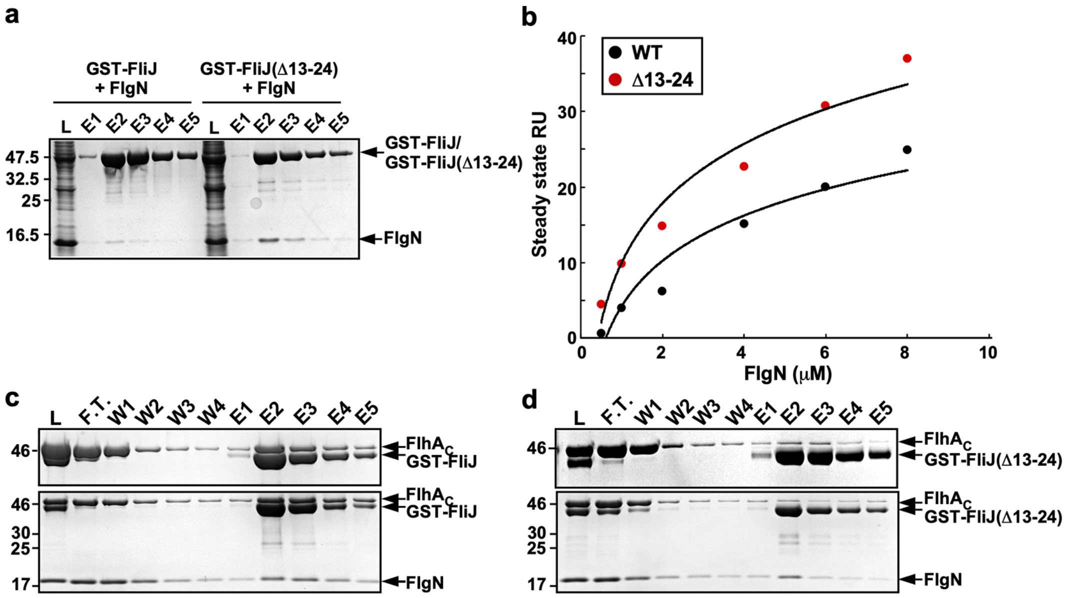

**Supplementary Fig. 4. Effect of a deletion of residues 13–24 of FliJ on interactions of FliJ with FlgN and FlhA$_C$. (a)** Interaction between FliJ and FlgN. Mixtures of cell lysates (indicates as L) prepared from SJW1368 (Δ*flhDC-cheW*) cells expressing either GST-FliJ or GST-FliJ(Δ13–24) with those from BL21(DE3) Star producing His-FlgN were loaded onto a GST column. After extensive washing, proteins were eluted with a buffer containing 10 mM reduced glutathione. Eluted fractions were analyzed by SDS-PAGE with CBB staining. **(b)** Measurements of the binding affinities of FliJ and FliJ(Δ13–24) for FlgN by SPR. His-FlgN of various concentrations was flowed over the sensor surface with immobilized GST-FliJ or GST-FliJ(Δ13–24) in 10 mM HEPES pH 7.4, 0.15 M NaCl, 3 mM EDTA, 0.005% Surfactant P20 at a flow rate of 20 μl/min. All experiments were performed at 25ºC. The steady-state resonance units (RU) were plotted against FlgN concentrations. **(c, d)** Effect of FlgN on the FliJ–FlhA interaction. Purified His-FlhA$_C$ was mixed with purified GST-FliJ **(c)** or GST-FliJ(Δ13–24) **(d)** in the absence (upper panel) and presence (lower panel) of purified His-FlgN, and dialyzed overnight against PBS. Each mixture (L) was loaded onto a GST column. After washing with 10 ml PBS at a flow rate of about 0.5 ml/min, proteins were eluted with 10 mM reduced glutathione. Flow through fraction (F.T.), wash fractions (W), and elution fractions (E) were analyzed by SDS-PAGE with CBB staining.

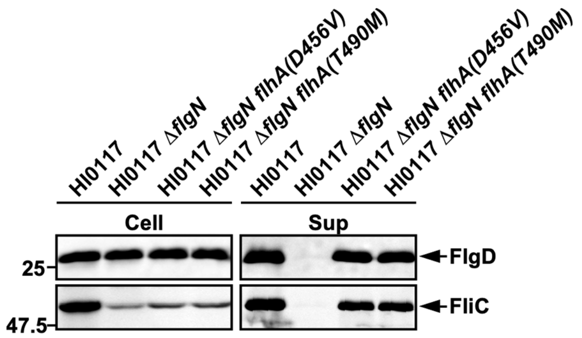

**Supplementary Fig. 5. Effect of the *flhA(D456V)* and *flhA(T490M)* mutations on flagellar protein export by the Δ*fliH–fliI flhB(P28T)* Δ*flgN* mutant.** Immunoblotting, using polyclonal anti-FlgD (1st row) or anti-FliC antibody (2nd row), of whole cell proteins (Cells) and culture supernatant fractions (Sup) prepared from the MMHI0117 [Δ*fliH–fliI flhB(P28T)*, indicated as HI0117], MM9002 [Δ*fliH–fliI flhB(P28T)* Δ*flgN*, indicated as HI0117 Δ*flgN*], HMM001 [Δ*fliH–fliI flhB(P28T)* Δ*flgN flhA(D456V)*, indicated as HI0117 Δ*flgN flhA(D456V)*] and HMM002 [Δ*fliH–fliI flhB(P28T)* Δ*flgN flhA(T490M)*, indicated as HI0117 Δ*flgN flhA(T490M)*].

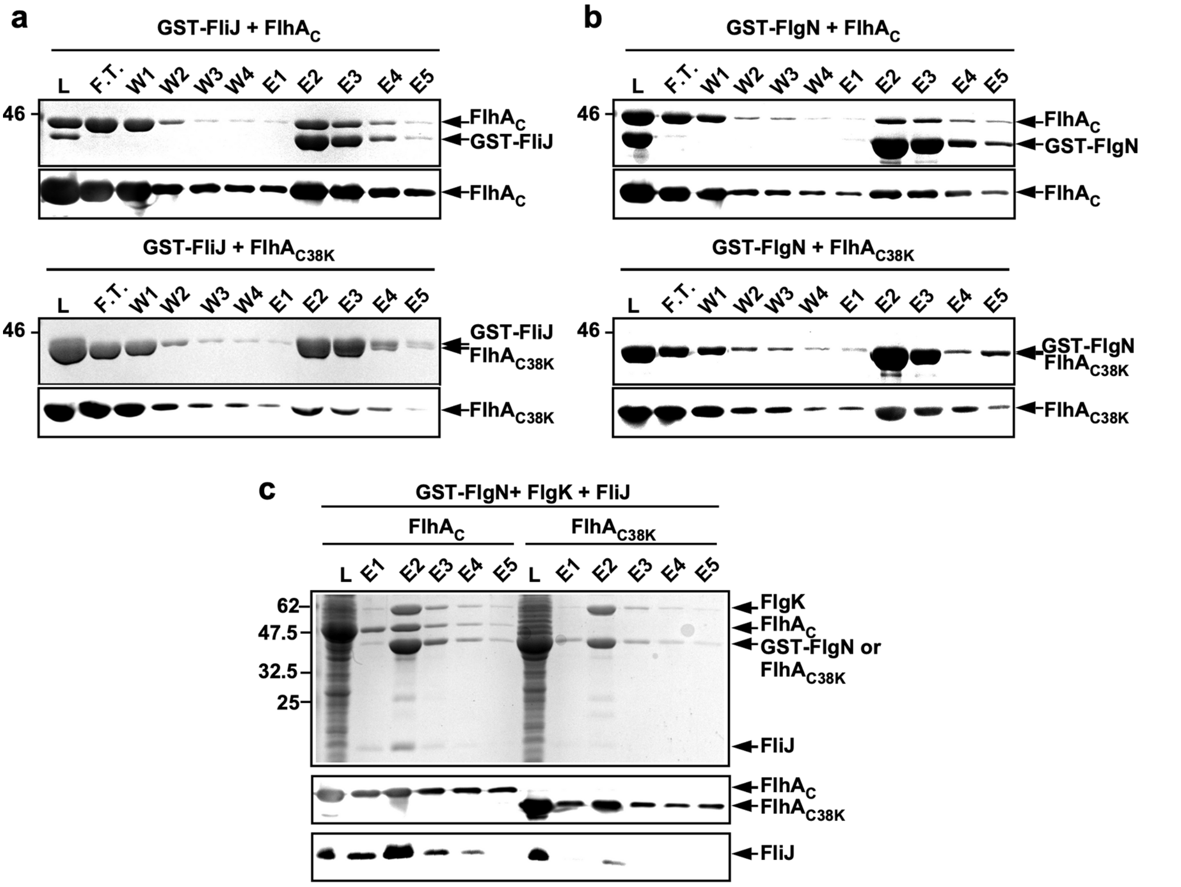

**Supplementary Fig. 6. Interactions of FlhA$_C$ with FliJ and FlgN. (a, b)** Effect of deletion of residues 328–351 of FlhA$_L$ on interactions of FlhA$_C$ with FliJ **(a)** and FlgN **(b)**. Purified His-FlhA$_C$ or His-FlhA$_{C38K}$ lacking residues 328–351 was mixed with purified GST-FliJ or GST-FlgN, followed by overnight dialysis against PBS at 4ºC. Each mixture (L) was loaded onto a GST column. After washing with 10 ml PBS at a flow rate of about 0.5 ml/min, proteins were eluted with 10 mM reduced glutathione. Flow through fraction (F.T.), wash fractions (W), and elution fractions (E) were subjected to SDS-PAGE, followed by both CBB staining (upper panels) and immunoblotting with polyclonal anti-FlhA$_C$ antibody. **(c)** Effect of deletion of residues 328–351 of FlhA$_C$ on the FlgN–FliJ interaction. Pull-down assays by GST affinity chromatography. Eluted fractions were analyzed by CBB staining (1st row) and immunoblotting with polyclonal anti-FlhA$_C$ (2nd row) or anti-FliJ (3rd row).

**Supplementary Table 1. Effect of $Na^+$ ions on average number and length of flagellar filaments produced by MM104-3 cells**

| Strain | NaCl (mM) | Fraction of flagellated cells (%) | Average number of flagella in flagellated cell (mean ± SD) | Average length of filament (μm) (mean ± SD) |
|---|---|---|---|---|
| SJW1103 | 0 | 100 (n = 152) | 2.7 ± 1.1 (n = 152) | 11.3 ± 2.1 (n = 50) |
| | 100 | 100 (n = 153) | 3.3 ± 1.5 (n = 153) | 12.9 ± 2.5 (n = 50) |
| MM104-3 | 0 | 61.5 (n = 192) | 1.3 ± 0.5 (n = 118) | 7.0 ± 2.9 (n = 50) |
| | 100 | 87.3 (n = 166) | 2.0 ± 1.0 (n = 145) | 10.7 ± 3.1 (n = 50) |

**Supplementary Table 2. Effects of gain-of-function mutations in FlhA and loss-of-function in FlgN on average filament number and length in flagellated MMHI0117 cells**

| | Fraction of flagellated cells (%) | Average filament number in flagellated cell (mean ± SD) | Average filament length (μm) (mean ± SD) |
|---|---|---|---|
| MMHI0117 (Δ*fliHI flhB**) | 78.2 (n = 340) | 1.6 ± 0.7 (n = 266) | 7.8 ± 2.5 (n = 50) |
| MMHIJ0117 (Δ*fliHIJ flhB**) | 13.5 (n = 423) | 1.1 ± 0.2 (n = 57) | 5.1 ± 2.2 (n = 50) |
| MMHIJ0117-2 [Δ*fliHIJ flhB* flhA(D456V)*] | 95.3 (n = 359) | 1.9 ± 0.8 (n = 342) | 5.7 ± 1.5 (n = 50) |
| MMHIJ0117-3 [Δ*fliHIJ flhB* flhA(T490M)*] | 99.2 (n = 376) | 2.1 ± 0.9 (n = 373) | 5.3 ± 1.5 (n = 50) |
| MM9002 (Δ*fliHI flhB**Δ*flgN*) | 0 (n = 266) | - | - |
| MM9004 (Δ*fliHIJ flhB**Δ*flgN*) | 0 (n = 329) | - | - |
| MM9004-2 [Δ*fliHIJ flhB**Δ*flgN flhA(D456V)*] | 20.8 (n = 355) | 1.0 ± 0.2 (n = 74) | - |
| MM9004-3 [Δ*fliHIJ flhB**Δ*flgN flhA(T490M)*] | 8.5 (n = 377) | 1.0 ± 0.0 (n = 32) | - |

### Table S3. Plasmids and *Salmonella* strains used in this study

| Strains/ Plasmids | Relevant characteristics | Source or reference |
|---|---|---|
| ***E. coli*** | | |
| BL21 (DE3) Star | Over-expression of proteins | Novagen |
| ***Salmonella*** | | |
| SJW1103 | Wild type for motility and chemotaxis | 1 |
| SJW1368 | Δ*cheW–flhD* | 2 |
| MM104-3 | *fliJ*(Δ13–24) *fliH*(Δ96–97) | 3 |
| MM9001 | Δ*flgN*::*tetRA* | 4 |
| MM9002 | Δ*fliH-fliI flhB(P28T)* Δ*flgN*::*tetRA* | 4 |
| MMHI0117 | Δ*fliH-fliI flhB(P28T)* | 5 |
| MMHI0117-2 | Δ*fliH-fliI flhB(P28T) flhA(D456V)* | 6 |
| MMHI0117-3 | Δ*fliH-fliI flhB(P28T) flhA(T490M)* | 6 |
| MMHIJ0117 | Δ*fliH-fliI-fliJ flhB(P28T)* | 3 |
| HMM001 | Δ*fliH-fliI flhB(P28T)* Δ*flgN*::*tetRA flhA(D456V)* | 4 |
| HMM002 | Δ*fliH-fliI flhB(P28T)* Δ*flgN*::*tetRA flhA(T490M)* | 4 |
| MM9003 | *fliJ*(Δ13–24) *fliH*(Δ96–97) Δ*flgN*::*tetRA* | This study |
| MM9003-2 | *fliJ*(Δ13–24) *fliH*(Δ96–97) Δ*flgN*::*tetRA flhA(D456V)* | This study |
| MM9003-3 | *fliJ*(Δ13–24) *fliH*(Δ96–97) Δ*flgN*::*tetRA flhA(T490M)* | This study |
| MM9003-4 | *fliJ*(Δ13–24) *fliH*(Δ96–97) Δ*flgN*::*tetRA fliI(L244R)* | This study |
| MM9003-5 | *fliJ*(Δ13-24) *fliH*(Δ96–97) Δ*flgN*::*tetRA flgM(S7ochre)* | This study |
| MM9004-6 | *fliJ*(Δ13-24) *fliH*(Δ96–97) Δ*flgN*::*tetRA* Δ*flgM* | This study |
| MMHIJ0117-2 | Δ*fliH-fliI-fliJ flhB(P28T) flhA(D456V)* | This study |
| MMHIJ0117-3 | Δ*fliH-fliI-fliJ flhB(P28T) flhA(T490M)* | This study |
| MM9004 | Δ*fliH-fliI-fliJ flhB(P28T)* Δ*flgN*::*tetRA* | This study |
| MM9004-2 | Δ*fliH-fliI-fliJ flhB(P28T)* Δ*flgN*::*tetRA flhA(D456V)* | This study |
| MM9004-3 | Δ*fliH-fliI-fliJ flhB(P28T)* Δ*flgN*::*tetRA flhA(T490M)* | This study |
| **Plasmids** | | |
| pTrc99AFF4 | Expression vector | 7 |
| pGEX-6p-1 | Expression vector | GE Healthcare |
| pMM104 | pET19b/ His-$FlhA_{C}$ | 8 |
| pMMHA001 | pET19b/ His-$FlhA_{C38K}$ | 9 |
| pMM406 | pTrc99A/ His-FliJ | 8 |
| pMKGK2 | pTrc99A/ FlgK | 10 |
| pMMGN101 | pGEX-6p-1/ GST-FlgN | 4 |
| pMMGN130 | pET15b/ His-FlgN | 11 |
| pMMGN140 | pTrc99AFF4/ FlgN | 11 |
| pMMJ1001 | pGEX-6p-1/ GST-FliJ | 12 |
| pMMJ1002 | pGEX-6p-1/ GST-FliJ(Δ13–24) | 3 |
